# Activating *RET* Mutations Promotes Osteoblastic Bone Metastases in Medullary Thyroid Cancer

**DOI:** 10.1101/2025.07.09.663851

**Authors:** Rozita Bagheri-Yarmand, Gabriel M. Pagnotti, Joseph L. Kidd, David Avila Ramirez, Trupti Trivedi, Leah Guerra, Zhongya Wang, Lucia Martinez Cruz, Jade M. Moehle, Yaashmin S. Jebaraj, Xinhai Wan, Jian H. Song, Roland Bassett, Sue-Hwa Lin, Mimi I. Hu, Steven I. Sherman, Marie-Claude Hofmann, Theresa A. Guise

## Abstract

Development of bone metastases increases mortality in patients with medullary thyroid cancer (MTC), with ∼50% survival at 5 years after diagnosis, but the underlying mechanisms are unknown. We show that patient-derived MTC cells (*RET*^C634W^ mutant TT cells and *RET*^M918T^ mutant MZCRC1 cells) promote an osteoblastic phenotype due to reduced bone resorption. Mechanistically, activated RET increases the expression of osteoprotegerin (OPG), an inhibitor of bone resorption, leading to decreased osteoclast differentiation. Furthermore, RET knockdown or pharmacological RET inhibition attenuates tumor burden and osteoblastic lesions in MTC-bearing femurs. Circulating levels of OPG were increased in the plasma of MTC patients who developed bone metastases and this was associated with poor overall survival. Patients who were treated with multi-kinase inhibitors have lower circulating levels of OPG. These novel findings identify a link between the RET signaling pathway and abnormal osteoblastic bone formation and suggest OPG as a potential biomarker of MTC bone metastases.

**Highlights:**

- Patient-derived MTC cells promote osteoblastic lesions in a mouse model.
- Activating the RET mutation promotes osteoprotegerin.
- Blocking RET kinase activity inhibits tumor growth and the osteoblastic lesion phenotype.
- High levels of circulating osteoprotegerin are associated with poor overall survival.

## INTRODUCTION

As of 2020, an estimated 558,260 people were living in the United States with a diagnosis of thyroid cancer and 38,857 new cases were estimated to be diagnosed that year (National Program of Cancer Registries, www.cdc.gov). Although the long-term survival outlook is favorable for most thyroid cancer patients, most individuals diagnosed with the medullary thyroid carcinoma (MTC) subtype will develop progressive tumor growth, aggressive infiltration, tumors with high vascularity, and prevalent metastatic disease. MTC uniquely arises from the parafollicular C cells of the thyroid glands that secrete calcitonin and carcinoembryonic antigen. At initial presentation, only about half (48%) of MTC patients have localized disease. Instead, 35% of patients with MTC have a tumor extending beyond the thyroid into the surrounding soft tissues or regional lymph node metastases, and 13-20% have distant metastases, typically to the lung, liver, or bones, which leads to a survival rate of ∼50% at 5 years after diagnosis [1, 2]. Bone metastases are associated with skeletal-related events (SRE), including pain, hypercalcemia, spinal cord compression, muscle weakness, and pathological fractures [1]. Although bone is a common site of metastasis for many cancers, effective therapies for bone metastases are still lacking[3]. Medullary thyroid cancer bone metastases are associated with osteoblastic, osteolytic, or mixed lesions[4]. However, the underlying mechanism(s) that generate these variable skeletal phenotypes in MTC remain unknown.

The human skeleton consists of two primary types of bone, cortical (also known as compact bone) and trabecular (also called spongy or cancellous) bone, and bone marrow. Bones are constantly being remodeled through the interactions of osteoblasts (OB), osteocytes (OCy), and osteoclasts (OC) [3]. Bone marrow mesenchymal stem cells (MSC) develop into pre-OB, then mature in bone-forming OB. OB produce collagen, osteocalcin, and osteopontin, which, together with hydroxyapatite and organic components, form the mineralized portion of bone. Mature OB line the mineralized surface of the bone. New bone is formed by deposition of protein matrix (or osteoid) by the OB, which become trapped within the structure and differentiate into OCy as the matrix mineralizes [5]. OC are large, multinucleated cells responsible for breaking down the mineralized bone tissue, which develops from the monocyte/macrophage lineage upon stimulation by receptor activator of nuclear factor kappa-β ligand (RANKL) as well as macrophage colony-stimulating factor (M-CSF) [6]. The maturation of MSC into OB involves activity of the transcription factors runt-related transcription factor 2 (RUNX2) and osterix (SP7), Wnt/β-catenin signaling, and bone morphogenetic protein 2 [7]. The maturation of OB is inhibited by expression of the Wnt inhibitors dickkopf 1 (DKK1) and sclerostin (SOST), as well as overexpression of cytokines such as fibroblast growth factor 23 (FGF23) [8–10]. Osteoprotegerin (OPG) is a decoy RANKL protein synthesized by OB and acts as a negative regulator of osteoclastogenesis [11]. Interaction of cancer cells with different niches may dictate the cellular fates and therapeutic responses of disseminated cancer cells and microscopic metastases.

The *RET (rearranged during transfection)* proto-oncogene encodes a 170 kDa transmembrane tyrosine kinase receptor. *RET* is the driver oncogene in ∼50% of MTC cases where a germline or somatic activating mutation results in ligand-independent constitutive activation of the receptor[12]. Of the 60 *RET*-activating mutations discovered to date, *RET* ^M918T^ is the most common and most aggressive mutation [13]. This mutation is located in the intracellular kinase domain of the receptor and results in a 10-fold increase in ATP-binding affinity, leading to a more stable receptor-complex and increase in auto-phosphorylation activity [14]. The *RET*^W634R^ mutation is located in the cysteine-rich domain of the RET extracellular domain that promotes receptor dimerization and constitutive activation of RET independent of ligand binding [15, 16]. To date, no studies have stratified MTC-skeletal related events and morphology based on the status of RET mutations.

Current treatment strategies for treating MTC metastatic bone disease include surgery with or without radiotherapy associated with non-selective multi-kinase inhibitors (vandetanib, cabozantinib) or selective RET inhibitors (selpercatinib) and anti-resorptive bone agents such as zoledronic acid or denosumab [17]. Cabozantinib is a receptor tyrosine kinase inhibitor with potent activity against multiple receptor tyrosine kinases, including VEGFR2, RET, and MET. Cabozantinib treatment affects the bone microenvironment, including reduced OC and increased OB numbers compared with control and elongation of the epiphyseal growth plate; in particular, a hypertrophic chondrocyte zone has been observed [18]. Selpercatinib is an ATP-competitive small molecule RET inhibitor that the US Food and Drug Administration approved for the treatment of lung cancer or MTC harboring RET alterations (mutations, fusions). In contrast to multi-kinase inhibitors, selpercatinib possesses selective, nanomolar potency against RET and a diverse set of RET mutations, including some acquired resistance mutations. Selpercatinib exhibits potent anti-tumor activity in patient-derived RET fusion-positive and RET-mutant mouse models [19]. Selpercatinib significantly improves progression-free survival more than cabozantinib and vandetanib in patients with progressive, advanced RET-mutant MTC who have not previously received treatment with a kinase inhibitor [20].

The optimal management of bone metastases in MTC is unknown. Hence, there is an urgent need to understand the interactions of MTC and the bone microenvironment to find biomarkers of aggressive disease and novel therapeutic targets [13, 21–23]. To our knowledge, the effect of RET-specific inhibitors on MTC bone metastasis-bearing models has not been reported. We propose the development of murine models of MTC tumor growth in bone and hypothesize that pharmacologic RET inhibition can effectively target MTC bone metastases to reduce tumor growth in bone, as well as improve patient morbidity and mortality.

Here, we established the first clinically relevant mouse model of bone metastases in MTC to better understand the interaction between MTC and the bone microenvironment and test RET inhibitors’ efficacy in treating MTC bone metastasis. We showed that *RET* mutations (C634W and M918T) promote a distinct osteoblastic phenotype in mice implanted with patient-derived MTC cells by increasing bone formation and decreasing bone resorption. The knockdown of RET in the MTC-*RET*^M918T^ model significantly reduced osteoblastic lesions, and the RET-specific inhibitor selpercatinib inhibited tumor growth in bone. Furthermore, ONC201, a small molecule inhibitor that we have previously shown to reduce RET at the protein level, inhibited tumor growth in bone. We showed that activated RET promotes OPG expression, which could be inhibited by RET inhibitors. Moreover, high OPG expression in MTC tumor tissues and plasma of MTC patients was associated with poor overall survival and may be considered as a biomarker of aggressive disease.

## RESULTS

### Patients-derived MTC cells with RET mutations (C634W and M918T) develop distinct osteoblastic bone metastases in mice

To investigate the tumor-bone interaction in MTC-bone metastasis, we injected patient-derived MTC-TT cells (RET-C634W mutation) or MTC-MZCRC1 cells (RET-M918T mutation) into the femur of SCID mice (NOD.Cg-Prkdcscid Il2rgtm1Wjl/SzJ) and allowed them to grow for 8 weeks. The incidence of tumor engraftment was 100 %.

Hematoxylin and eosin staining showed that TT cells induced woven bone lined by cuboidal OB within the bone marrow cavity (Figure 1A). Micro-computed tomography (micro-CT) reconstruction of the entire femur of mice injected with MTC-TT cells with the RET^C634W^ mutation showed a four-fold increase in trabecular bone volume, trabecular number, and a two-fold increase in trabecular thickness compared with non-tumor-bearing mice without significant changes in cortical parameters (Figure 1B). DXA analysis confirmed a two-to three-fold increase in bone mineral density (Figure 1C). Tartrate-resistant acid phosphatase (TRAP) staining was performed to detect OC. The number of TRAP-positive, multinuclear OC in the bone-tumor interface from mice injected with TT cells was significantly lower than in non-injected femur (*p*<0.01; Figure 1D). These results suggest that mice injected with MTC cells exhibited increased bone formation with less resorption. The number of active OB was decreased at the bone surface (Figure 1E), in contrast to the number of OCy per bone surface, which was significantly increased in the cortical region (*p* <0.01), suggesting that the mature OB may have been differentiated into OCy (Figure 1F). Von Kossa staining of the femurs of TT-injected mice showed a three-fold increase in mineralization of the trabecular region without significant increase in the cortical area (Figure 1G).

**Figure 1.**
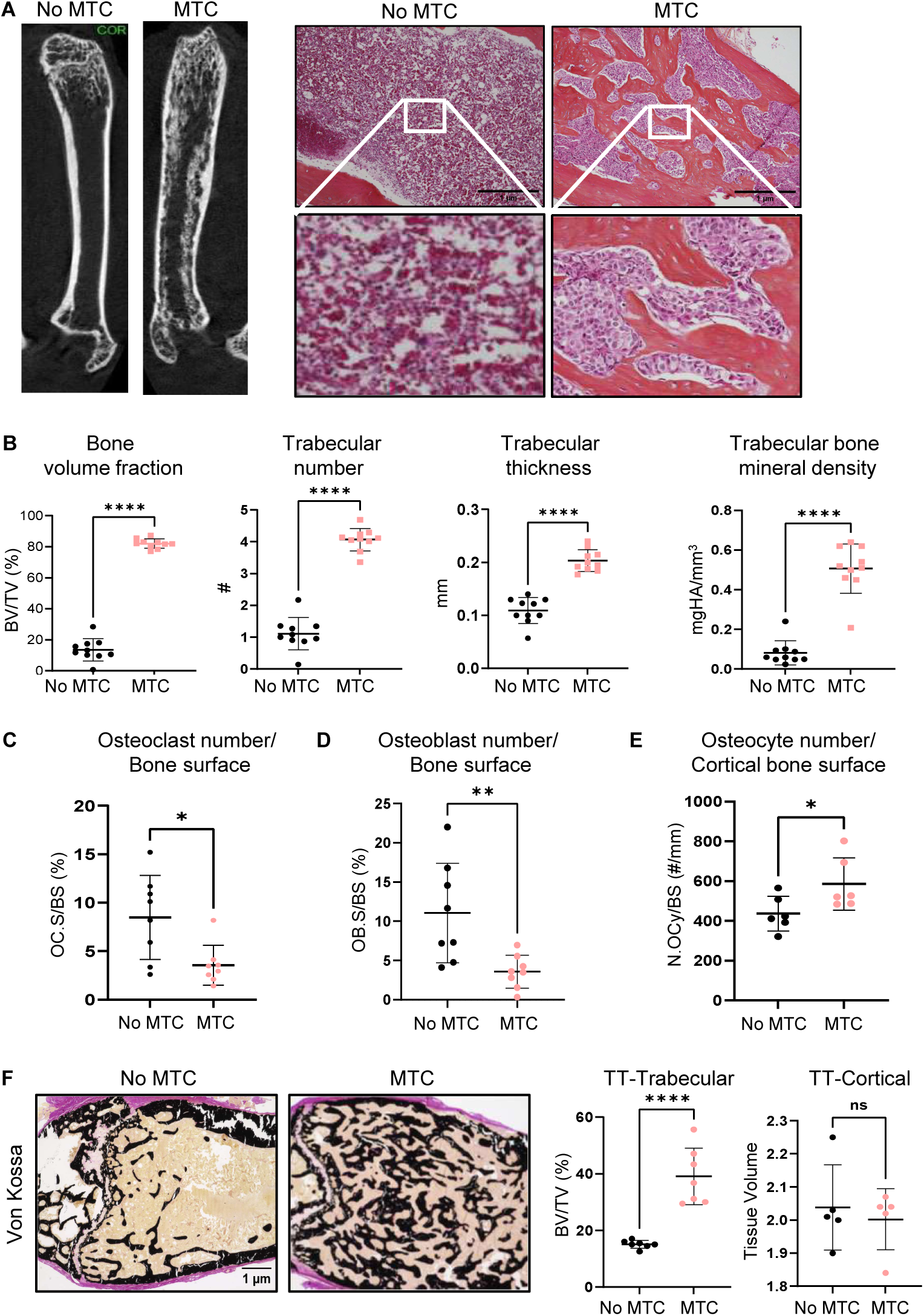
Patient-derived medullary thyroid cancer-TT cancer cells enhance osteoblast (OB)- mediated bone formation and inhibit osteoclast (OC)-mediated bone resorption in an *in vivo* murine model. **A)** Representative 3D reconstructions micro-computed tomography (micro-CT) - images and hematoxylin and eosin histologic analysis of MTC-TT tumor-bearing male NOD/SCID mouse femoral bone 8 weeks after inoculation. **B)** Micro-CT quantification of bone microarchitecture parameters. BMD, bone mineral density; BV/TV, trabecular volume per total bone volume; Tb.N, average trabeculae number; Tb.Th, average trabeculae thickness (n=10). Bone histomorphometry analysis (*Bioquant; OSTEO v.20.1*) of **C)** percentage of bone surface (BS) covered with OC (OC.S/BS); **D)** percentage of total bone surface lined with OB (OB.S/BS); and **E)** the number of osteocytes (OCy) per cortical bone surface (OCy #/BS) in MTC-TT bearing mouse femurs. **F)** Von Kossa staining and quantification in trabecular and cortical regions showing increased mineralization. Data are represented as individual values and averages with SD bars (n=10 mice per group). Two-tailed Student’s t test determined *p-values*. *, *p* <0.05; **, *p* <0.01; ***, *p*<0.001; and ****, *p*<0.0001.

SCID mice inoculated with MZCRC1 cells harboring the RET^M918T^ mutation showed increased trabecular bone volume, cortical thickness and porosity with reduced medullary area compared with non-tumor-bearing mice in micro-CT analysis (Figure 2A). Von Kossa staining of the MZCRC1-injected femurs showed a threefold increase in mineralization in the trabecular region. Also, a significant increase was observed in the cortical area (p<0.01; Figure 2B). Calcein double labeling showed an increase in bone formation rate across the cortical metaphysis (Figure 2C).

**Figure 2.**
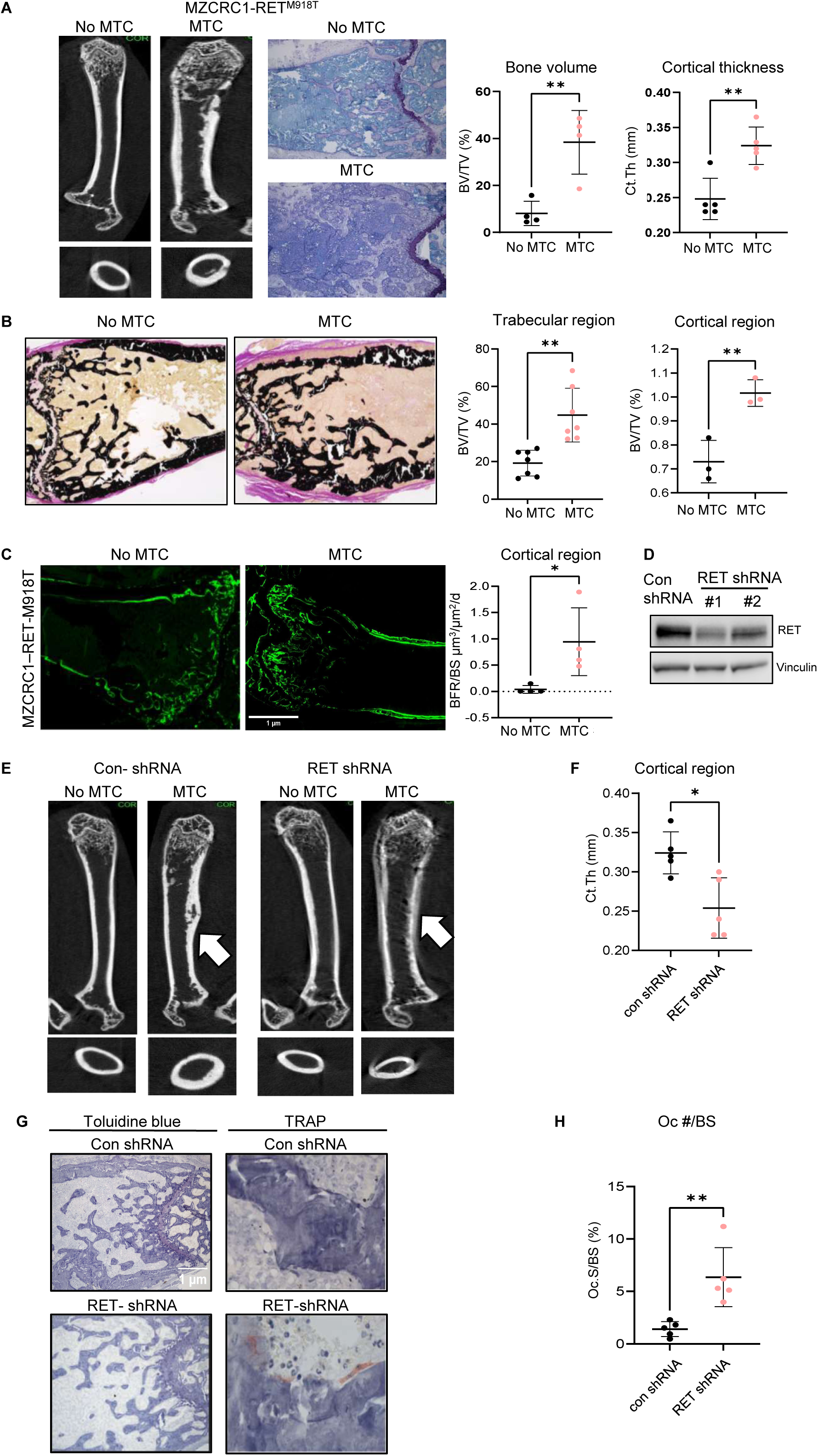
Patient-derived medullary thyroid cancer (MTC)-MZCRC1 cancer cells enhance osteoblast (OB)-mediated bone formation in cortical regions. **A)** Representative 3D reconstructions micro-computed tomography (micro-CT) images and Toluidine blue staining of MTC-MZCRC1 tumor-bearing male NOD/SCID mouse femoral bone 8 weeks after inoculation. Micro-CT quantification of bone microarchitecture parameters is also shown. Trabecular BV/TV, bone volume fraction; Tb.Th, average trabecular thickness. **B**) Von Kossa staining and quantification in trabecular and cortical regions showing increased mineralization. **C)** Bone formation rate (BFR/BS) was measured by calcein injection on undecalcified bone sections. **D)** RET knockdown in MTC-MZCRC1. MZCRC1 cells stably expressing RET-specific shRNA were generated by lentiviral shRNA transduction, and decreased RET protein levels are shown by Western blot analysis in 2 stable cell lines. **E)** RET knockdown in MTC-MZCRC1 cells reduced cortical thickness and OC number per bone surface. The representative micro-CT images of bone formation in trabecular and cortical regions in the femurs of SCID mice injected with MTC-MZCRC1 cells are shown. **F)** Toluidine blue and TRAP staining of femoral regions in control shRNA and RET knockdown and number of osteoclast (OC) per bone surface. Data are represented as individual means ± SD (n=10 mice per group). Two-tailed Student’s t test determined *p-values*. *, *p* <0.05; **, *p* <0.01; ***, *p* <0.001; and ****, *p* <0.0001.

### RET knockdown in MTC-MZCRC1 cells decreases cortical thickness and porosity

To examine the role of RET^M918T^ in the development of osteoblastic lesions, we knocked down RET expression in MTC-MZCRC1 cells using RET short hairpin RNA (shRNA). Cells with non-targeted shRNAs served as controls. Western blot analysis showed that RET protein levels in RET-shRNA cells were decreased by 40-60% (Figure 2D). We injected two stable MZCRC1-RET knockdown cell lines into the right femurs of SCID mice, which significantly reduced cortical thickness and porosity (Figure 2E, F). TRAP staining revealed that OC numbers were significantly increased in RET-shRNA tumors compared with tumors with control-shRNA (*p*<0.0001; Figure 2G, H). These results indicate that MTC-induced osteoblastic lesions are mediated by RET.

### Conditioned media of MTC increases OB differentiation

In order to understand the mechanisms of tumor:bone interactions in MTC, we performed *in vitro* experiments with conditioned media. Since OB are the primary bone-forming cells, we investigated the effect of MTC-conditioned media (MTC-CM) on mouse calvarial OB (MCO) differentiation. MCO were isolated from the calvaria of newborn C57Bl6 mice and cultured with MTC-CM in media supplemented with ascorbic acid and β-glycerophosphate for 2 weeks. Treatment of MCO with TT-CM increased OB differentiation, as assessed by alkaline phosphatase staining (Figure 3A). CM from TT cells with RET knockdown inhibited TT-mediated OB differentiation (Figure 3A). We also found that MCO culture with TT-CM increases mineralization, as evidenced by alizarin red-S-stained calcified nodules, and this effect was abrogated upon RET knockdown (Figure 3A, B). We also performed co-culture studies with TT-CM and MZCRC1-CM using bone marrow MSC isolated from 7-week-old C57Bl6 mice. The results showed increased OB differentiation in culture with MTC-CM (Figure 3C). Furthermore, quantitative reverse-transcriptase polymerase chain reaction (qRT-PCR) analysis showed that treatment of MCO with TT-CM increased mRNA levels of the osteogenic genes, including ALP, OCN, SP7, and OPG, whereas the expression of SOST and RANKL were decreased (Figure 3D). Together, these results suggest that MTC cells with activated RET kinase acts on pre-OB to promote OB differentiation and mineralization.

**Figure 3.**
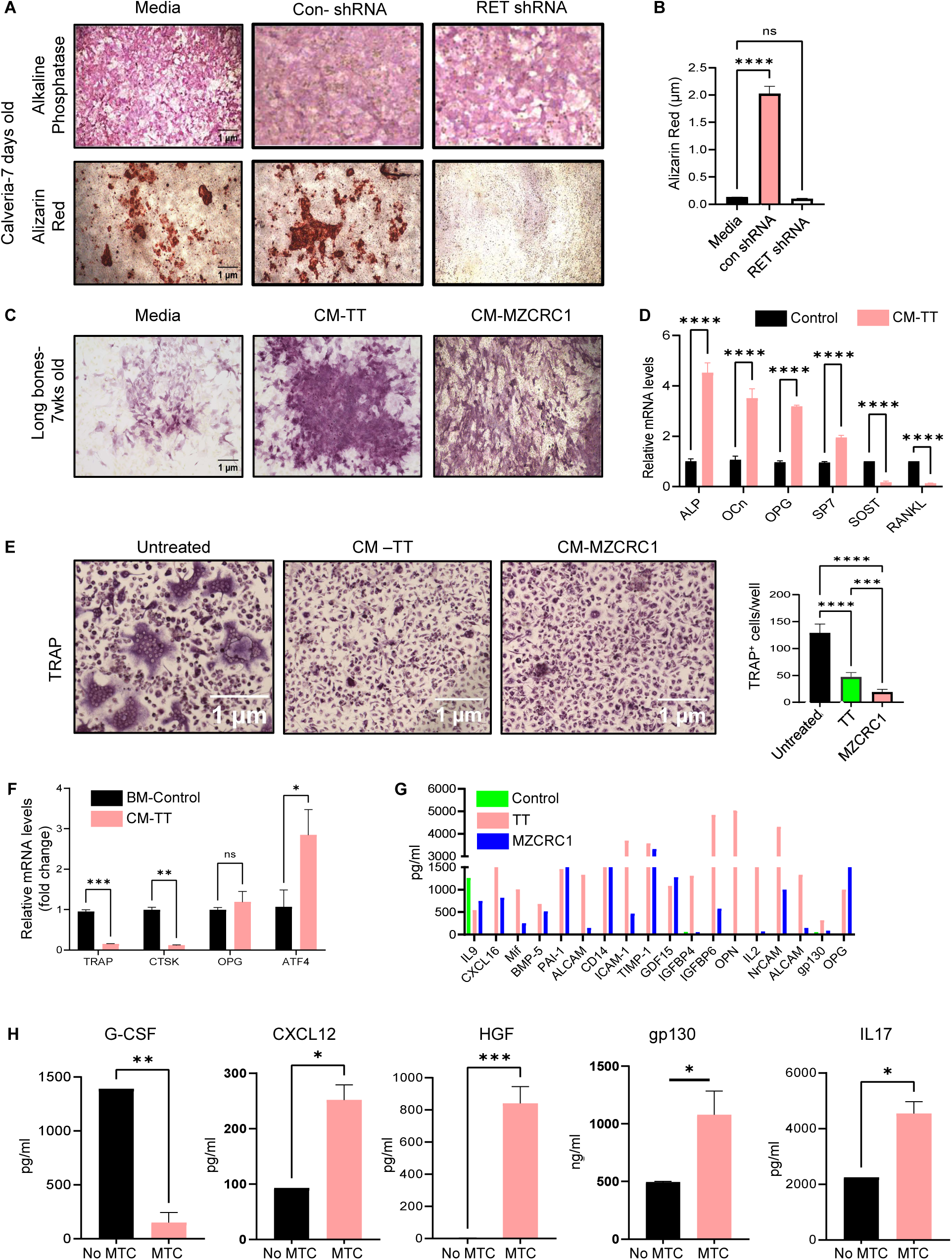
Medullary thyroid cancer (MTC)-secreted factors promote osteoblast (OB) differentiation. **A)** Pre-OB were isolated from the calvaria of newborn mice and cultured with conditioned media (CM) from TT or TT-RET shRNA in differentiation media for 2 weeks and stained for alkaline phosphatase (ALP) and alizarin red. Representative images of ALP and alizarin red are shown. **B)** Quantification of alizarin red staining with cetylpyridinium chloride extraction. **C)** Pre-OB were isolated from the long bones of 7-week-old mice, cultured with CM from TT or MZCRC1 cells in differentiation media for 2 weeks, and stained for ALP. **D)** Total RNA was harvested, real-time quantitative reverse-transcriptase PCR (qRT-PCR) for osteogenic genes (ALP; OCN (osteocalcin); OPG (osteoprotegerin), (Sp7 (osterix); SOST, RANKL) was performed, and gene expression was normalized to that of β-actin. **E)** MTC-secreted factors inhibit osteoclastogenesis. Bone marrow-derived macrophages from C57BL/6J mice were induced to differentiate toward osteocalcin (OC) with M-CSF and RANKL in the presence or absence of CM from TT and MZCRC1 cells. After 7 days of culture, the cells were stained for TRAP and photographed to observe the multinucleated TRAP+ cells (fused cells with more than 3 nuclei) known as OC. Quantification of multinucleated TRAP+ cells in different experimental groups is shown. **F)** Quantification of OC differentiation markers by qRT-PCR (CTSK (cathepsin K), RANKL, OPG, and ATF4). **G)** Quantification of cytokine and chemokine protein concentration in CM from TT and MZCRC1 cells (*RayBiotech*). **H)** Mouse cytokine array from the serum of mice inoculated with TT cells (MTC) and control non-injected (no MTC) in bone after 8 weeks. Data are represented as means ± SD (n=10 mice per group). Two-tailed Student’s t test determined *p-values*. *, *p* <0.05; **, *p* <0.01; ***, *p* <0.001; and ****, *p* <0.0001.

### MTC-CM inhibits osteoclastogenesis

Because MTC cells reduced OC numbers in the bone microenvironment in mice, we examined whether CM of both MTC cells (TT and MZCRC1 cells) regulates OC formation. We isolated bone marrow monocytes from 7-week-old C57BL/6J mice and cultured the cells in the presence of M-CSF and RANKL to stimulate OC formation. Treatment with MTC-CMs resulted in a significant decrease in TRAP-positive multinucleated cells (Figure 3E). Real-time qRT-PCR confirmed the decreased expression of osteoclastogenesis marker TRAP and CSTK and increased expression of ATF4 which is a maker of primary bone marrow cells when co-cultured with MTC-CM (Figure 3F). These results indicated that MTC-CM contains cytokines that suppress osteoclastogenesis.

### MTC secretes cytokines that inhibit osteoclastogenesis and promote bone formation

Dynamic regulation of osteoclastogenesis and anti-osteoclastogenesis cytokines is essential in maintaining the balance between bone-resorbing OC and bone-forming OB [24]. It has been reported that soluble factors and cytokines secreted from tumor cells are directly disturbing bone homeostasis. We tested whether MTC cells release cytokines that change bone homeostasis. To this end, bone marrow of TT and MZCRC1 cells was collected and tested by cytokine array (*RayBiotech*). This analysis showed that the expression of several pro-inflammatory cytokines was increased, including IL9, CXCL16, CD14, IGFBP6, osteopontin, IL2, GDF15, and OPG in TT-CM and MZCRC1-CM compared with the control medium (Figure 3G).

OPG inhibits OC differentiation and is a decoy receptor of RANKL [11], suggesting that high OPG secretion in the MTC-CM may cause inhibition of osteoclastogenesis. We further tested whether MTC-mediated change in the bone microenvironment leads to changes in the cytokine profile in the serum. To this end, the serum of mice injected with TT/MZCRC1 cells in the femur was analyzed for cytokine profiling by ELISA immunoassays (*RayBiotech*). We observed increased cytokine/growth factor levels in the serum of mice injected with TT cells in bone, including CXCL12 (SDF-1a), IL17B, and HGF. The secretion of granulocyte-macrophage-colony-stimulating factor (GM-CSF) in the serum was decreased by 10-fold (Figure 3H). GM-CSF has been shown to increase OC formation and activity [25], and gp130 promotes bone formation by enhancing the differentiation of OB progenitors [26]. CXCL12 promotes the homing of cancer cells to the bone surface and cancer cell survival and has been shown to mediate breast cancer growth and dissemination. [27, 28]. IL17B stimulates OB differentiation and mineralization [29]. These findings suggest that MTC cells secrete cytokines to affect bone turnover.

### Activated RET mutations regulate OPG expression

Because we observed high levels of OPG in MTC-CM and downregulation of RET led to increased OC in the MTC tumors, we hypothesized that RET mutations upregulate OPG expression. To test our hypothesis, we examined OPG expression in MTC-RET knockdown cells and found that knockdown of RET in TT and MZCRC1 cells reduced OPG expression (Figure 4A, B). We then examined whether RET-specific inhibitors selpercatinib or pralsetinib inhibit OPG protein levels in MTC. As expected, both selpercatinib and pralsetinib inhibited RET autophosphorylation and OPG protein levels without changing OPG mRNA levels (Figure 4C, D). Next, we examined the effect of selpercatinib on OPG secretion. We observed that the levels of OPG secretion in MTC-CM were decreased in selpercatinib-treated cells by ELISA (Figure 4E). Previously, we reported that the small molecule ONC201 reduces RET expression in MTC *in vitro* and inhibits tumor growth *in vivo* [30]. Thus, we examined the effect of ONC201 on RET and OPG expression in MTC. We observed that treatment with ONC201 decreased RET protein in a dose-dependent manner and reduced OPG at the protein and mRNA levels (Figure 4F, G). These results suggest that activated RET enhances OPG protein secretion in MTC-CM. Knockdown of RET or pharmacological inhibition of RET activity reduced OPG levels.

**Figure 4.**
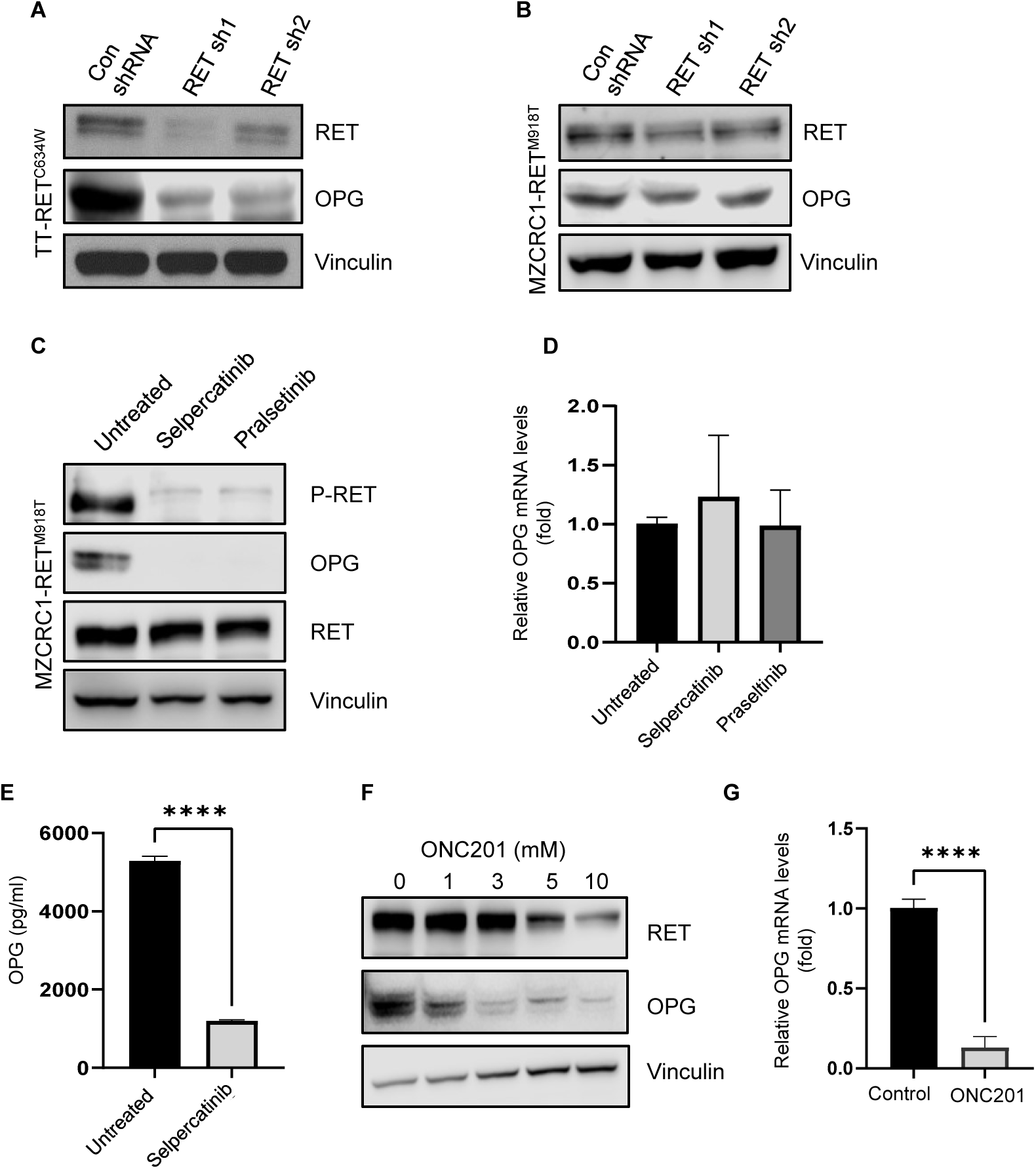
Osteoprotegerin (OPG) protein levels are decreased in RET knockdown MTC cells**. A)** Western blot analysis showing RET and OPG protein levels in stable RET knockdown by shRNA lentivirus in TT cells (stable clone #1 and #2). **B)** Western blot analysis showing RET and OPG protein levels in stable RET knockdown by shRNA lentivirus in MZCRC1 cells (stable clone #1 and #2). Vinculin served as a loading control. **C)** RET-specific inhibitors selpercatinib and pralsetinib inhibit RET phosphorylation and OPG protein levels. MZCRC1 cells were treated with selpercatinib or pralsetinib (5 μM) for 48 hr and subjected to Western blot analysis with the indicated antibodies. **D)** Real-time PCR quantitative PCR (qRT-PCT) showing mRNA levels of OPG in MZCRC1 cells treated with selpercatinib and praseltinib. Values were normalized against GAPDH mRNA levels. **E)** Selpercatinib decreased OPG secretion by MTC cells. MZCRC1 cells were treated with selpercatinib (5 μM) for 48 hr, and conditioned media (CM) was collected after 24 hr in media containing 0.1% serum and then analyzed by ELISA. **F)** MZCRC1 cells were treated with small molecule ONC201 for 72 hr with increasing indicated concentration and subjected to Western blot analysis with the indicated antibodies. **G)** ONC201 decreases OPG expression by MTC cells. MZCRC1 cells were treated with ONC201 (5 μM) for 48 hours, and isolated mRNA was analyzed by real-time qRT-PCR. Data are represented as averages of triplicates ± SD from two independent experiments. The two-tailed Student’s t-test determined *p-values*. *, *p* <0.05; **, *p* <0.01; ***, *p* <0.001; and ****, *p* <0.0001.

### RET-specific inhibitor selpercatinib inhibits tumor growth in mice

Selpercatinib has been approved by the US Food and Drug Administration for adult and pediatric patients (≥2 years of age) with advanced or metastatic MTC with a RET mutation, exhibiting a superior progression-free-survival compared with cabozantinib or vandetanib [20]. To investigate the effect of selpercatinib on tumor growth in bone, we inoculated TT and MZCRC1 cells (5.0 × 10^5^ per injection) into the right femur of male SCID mice. The left femur served as control. After 6 weeks, the presence of MTC tumors in bone was verified by Xray, and the daily selpercatinib treatment (30 mg/kg per day) was initiated. Treatment continued for 3 weeks (Figure 5A). No overt toxicities (weight loss, lethargy, or hunched posture) were noted in selpercatinib-treated mice (Supplemental Figure S1). Micro-CT analyses were performed in trabecular and cortical regions of the femur in TT- and MZCRC1-inoculated femurs and controls in untreated and treated mice. We observed a significant increase in BV/TV in TT-inoculated femurs compared with controls in the trabecular region but not cortical regions of the distal femoral metaphysis, and treatment with selpercatinib did not significantly change BV/TV compared with control mice (Figure 5B, C). Analysis of MZCRC1-inoculated femurs showed increased cortical thickness and bone volume, which significantly decreased with selpercatinib treatment (Figure 5D, E). Furthermore, selpercatinib inhibited tumor growth by 50% in both TT- and MZCRC1-inoculated femurs (Figure 5F, G). The number of OC per bone surface (N.OC/BS) was increased in selpercatinib-treated mice in both TT- and MZCRC1-inoculated femurs compared with control mice (Figure 5H). The number of OCy per bone surface (N.OCy/BS) was decreased in cortical regions of mice treated with selpercatinib compared with controls in both TT- and MZCRC1-inoculated femurs (Figure 5I). These results suggest that selpercatinib inhibits tumor growth and attenuates MTC-induced osteoblastic bone lesions.

**Figure 5.**
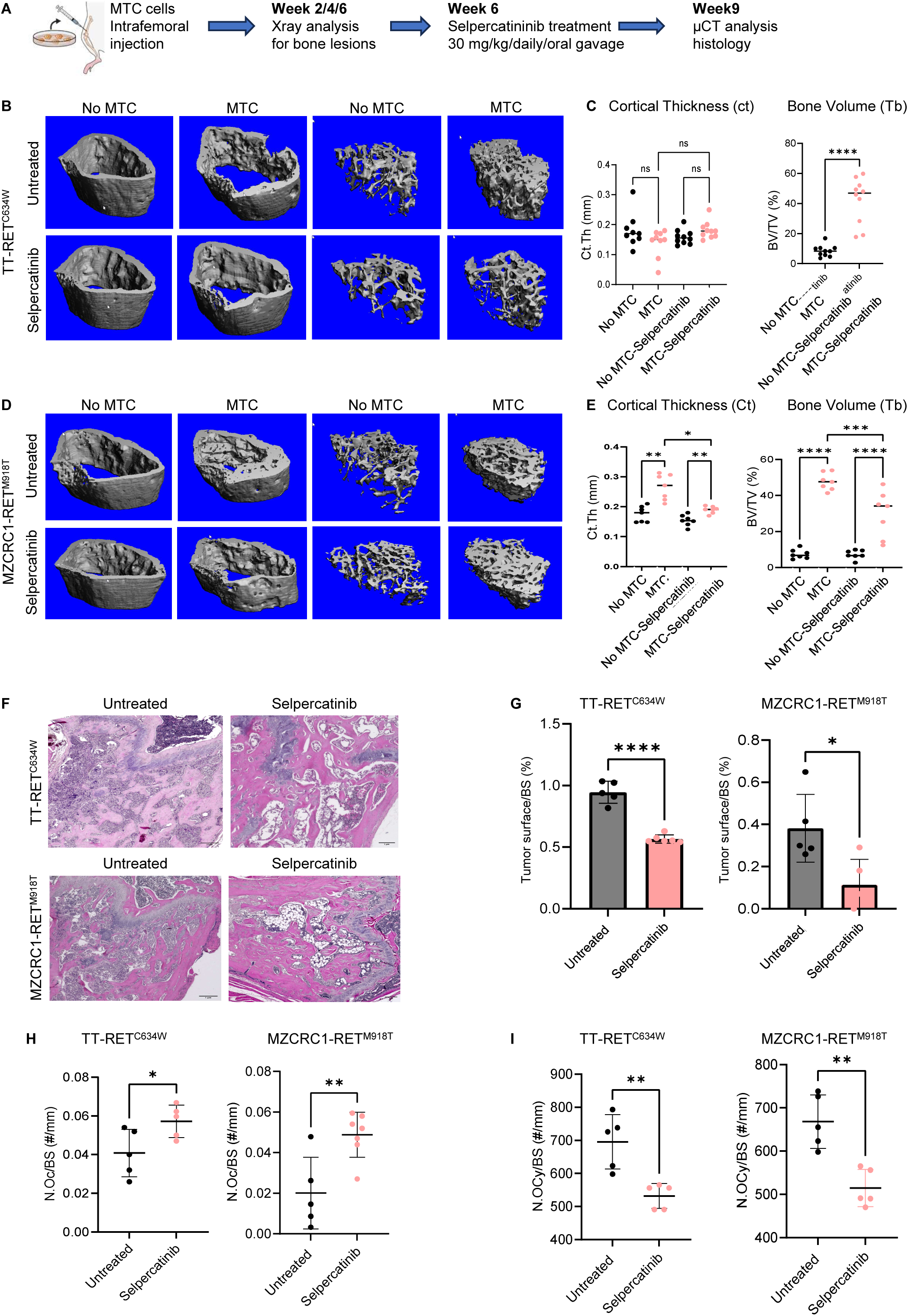
Treatment with selpercatinib attenuates osteoblastic lesions and reduces tumor growth. **A**) Schematic workflow of the *in vivo* mouse studies and examinations. **B, C**) Representative 3D reconstructed micro-computed tomography (micro-CT) images of medullary thyroid cancer (MTC)-TT and MZCRC1 cell tumor-bearing male NOD/SCID mouse femoral bone 3 weeks after treatment. **D)** Micro-CT quantification of bone microarchitecture parameters. BV/TV, trabecular bone volume fraction; Tb.Th, average trabeculae thickness. **E)** Hematoxylin and eosin staining of TT- and MZCRC-bearing femurs treated with selpercatinib. **F)** Quantification of tumor surface per bone surface of TT- and MZCRc1-bearing tumors treated with selpercatinib (*OSTEO v20.1, Bioquant Imaging Analysis*). Scale bar = 1 µm. G) Quantification of osteoclast (OC) per bone surface of TT- and MZCRC1-bearing femur. Data are represented as means ± SD. Two-tailed Student’s t-test determined *p-values*. *, *p* <0.05; **, *p* <0.01; ***, *p* <0.001; and ****, *p*<0.0001.

### ONC201 inhibits tumor growth in bone and mitigates tumor-induced bone disease

Based on previous studies, we hypothesized that inhibiting RET by ONC201 can inhibit tumor growth and reduce MTC-induced osteoblastic lesions. To assess the anti-tumor activity of ONC201 in bone, we inoculated MTC-TT cells (5.0×10^5^ cells per injection) into the right femur of 6-week-old male NGS/SCID mice (n=15). After 4 weeks, the mice were randomized into two groups based on Xray images; one group was an untreated control group and the other received ONC201 via oral gavage. Mice were treated once per week (120 mg/kg) for 6 weeks. Micro-CT analysis showed that bone volume fraction, trabecular thickness, and trabecular number per bone surface were reduced in mice treated with ONC201 (Figure 6A, B). In contrast, cortical thickness was lower in MTC-bearing femurs, and the cortical thickness increased with ONC201 treatment (Figure 6C). Histomorphometric analyses was decreased that ONC201 increased the number of OB at the bone surface compared with the vehicle (Figure 6E). Ki67 staining, a marker for cell proliferation, was decreased in tumor cells in mice treated with ONC201 (Figure 6F, G). Furthermore, treatment with ONC201 significantly decreased tumor volume (Figure 6H). Immunohistochemical staining showed decreased RET and OPG expression in ONC201-treated mice (Figure 6I, J, K). These results suggest that ONC201 treatment significantly reduced tumor growth and MTC-induced osteoblastic bone lesions.

**Figure 6.**
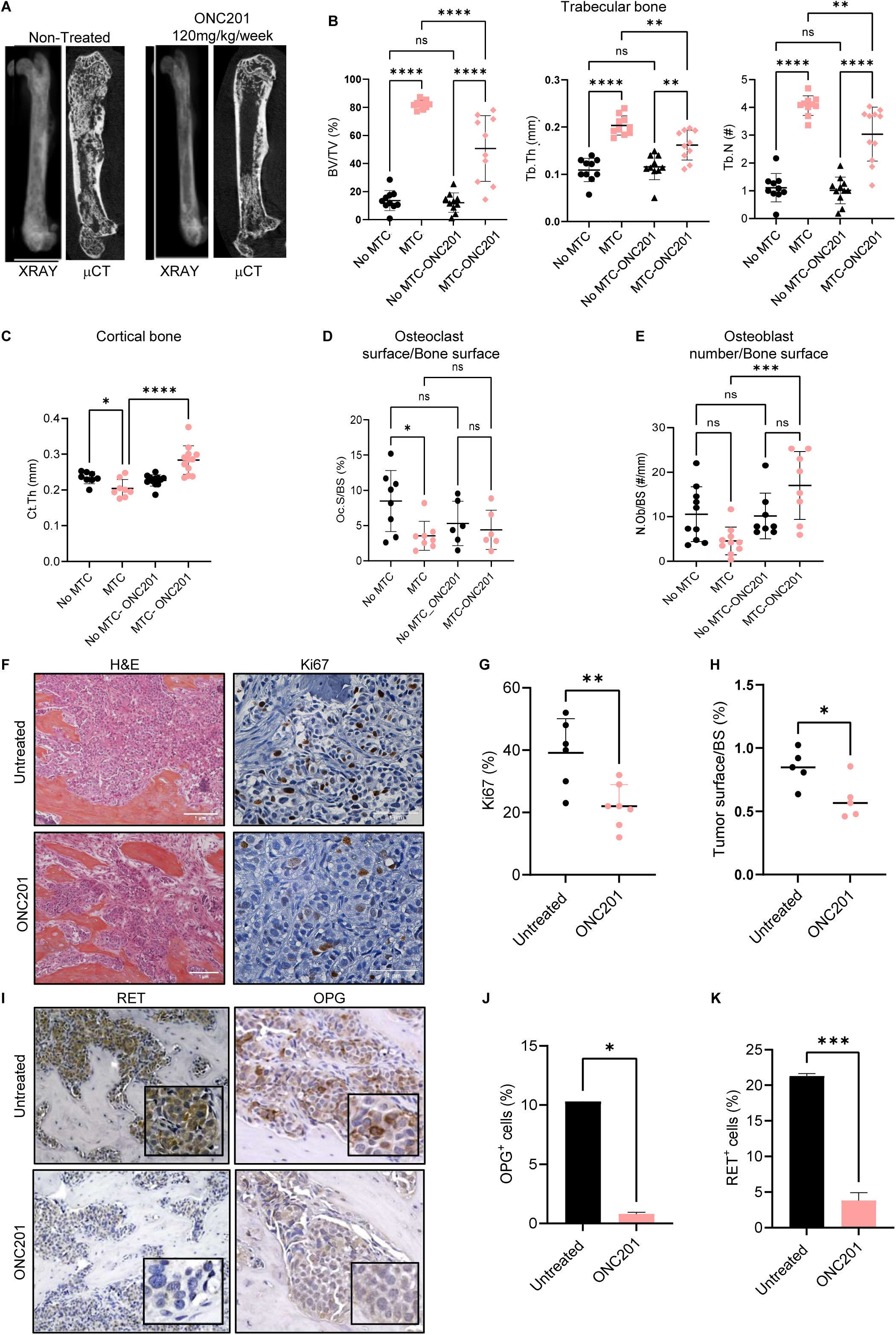
Treatment with the small molecule ONC201 attenuates osteoblastic lesions and reduces tumor growth. **A)** TT cells were injected into SCID mouse femurs, and 4 weeks after tumor were confirmed by X-ray, mice were treated with ONC201 at 120 mg/kg/once per week for 6 weeks. Representative X-ray and micro-computed tomography (micro-CT) images are shown in both control and treated mice **B)** Quantification of the trabecular region in medullary thyroid cancer (MTC)-bearing femurs showing the percentage of bone volume fraction (BV/TV), trabecular thickness, and trabecular number. **C)** Quantification of the cortical region in MTC-bearing femurs. **D)** Quantification of osteoclast (OC) surface per bone surface (*OSTEO v20.1, Bioquant Imaging Analysis*). **E)** Quantification of osteoblast (OB) number per bone surface. **F, G)** Representative immunohistochemical images of Ki67, a proliferation marker, and quantification of Ki67+ levels in MTC-TT-bearing femurs. **H**) Quantification of tumor surface per bone surface. **I, J, K**) Representative immunohistochemical images of RET and OPG and quantification in MTC-TT bearing femurs in control mice and mice treated with ONC201. Scale bar = 1 μm. Data are presented as means ± SD (n = 10 mice/group) Student’s *t-*test or ordinary one-way ANOVA determined *p-values*. *, *p* <0.05; **, *p* <0.01; ***, *p* <0.001; and ****, *p* <0.0001.

### Circulating OPG is a biomarker of MTC bone metastasis

To elucidate the clinical relevance of OPG in MTC bone metastases, we examined circulating OPG levels in 80 plasma samples and OPG protein expression in 20 MTC tumor tissue samples from 80 patients with MTC using immunohistochemistry (Supplemental Table S1). The median age at diagnosis was 63 years (range 28-83 years). Thirty-four of the 80 patients were male (42%), and 46 were female (58%). Fifty-four patients (68%) had stage IV disease. Forty-three patients (54%) presented with bone metastasis. All patients had non-hereditary(sporadic) disease, clinically verified through genetic testing. Fifty-two patients (65%) showed a somatic *RET^M918T^* mutation. Thirty-five patients were treated with the multi-kinase inhibitor vandetanib, cabozantinib, or both before blood collection.

To investigate OPG as a potential predictor of risk for bone metastasis development, we examined OPG protein levels in MTC tumor tissues by immunohistochemistry (Supplemental Table S2). We found that MTC patients who developed bone metastases had higher levels of OPG than patients without bone metastases (median: 1510 vs. 1097, *p*=0.002) (Figure 7A, B). Next, we examined circulating OPG levels using ELISA in the plasma samples from each of the 80 patients. The OPG plasma levels were significantly higher in patients with RET^M918T^ mutation who developed bone metastases than patients without RET^M918T^ mutations or patients with no metastases harboring RET^M918T^ mutation (Figure 7C). Moreover, patients treated with multi-kinase inhibitors before blood collection had lower levels of OPG in their plasma (Figure 7D). An OPG plasma ≤1100 pg/mL was considered low. Kaplan-Meier analysis showed that patients who developed bone metastases had a median overall survival of 181.4 months (95% CI 155.7 – 417.6), which was shorter than that of patients with no metastases whose median OS was not estimable (Figure 7E). The median overall survival was 331.6 months (95% CI 93.8 months to not reached; log-rank *p*=0.07) in patients with bone metastases and harboring RET^M918T^ mutation but was not yet reached in patients who did not develop bone metastases with RET^M918T^ mutation (Figure 7E). We found that MTC patients with high levels of OPG in their plasma have markedly shorter overall survival than patients with low OPG plasma levels (HR: 2.73, 95% CI 1.01 7.44; *p*=0.040) (Figure 7F, Supplemental Table S3). The median overall survival was 417.6 months in patients with high OPG plasma levels and was not reached in patients with low OPG plasma levels (Figure 7F).

**Figure 7.**
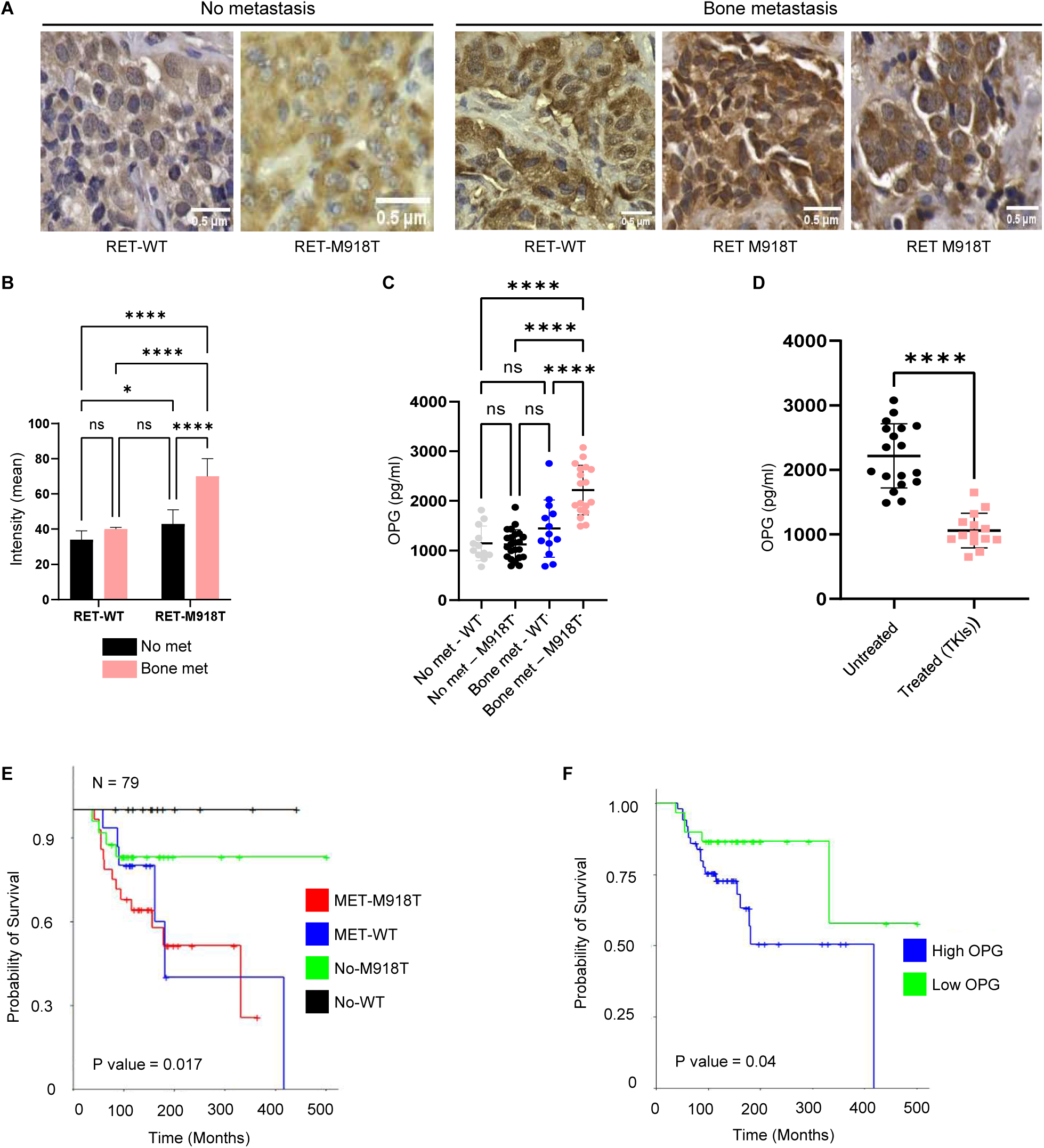
Osteoprotegerin (OPG) is a potential biomarker of bone metastases in medullary thyroid cancer (MTC). **A)** Representative immunohistochemical images of OPG in tumor samples from patients with primary sporadic MTC with or without RET mutations who did or did not develop bone metastases. Scale bars = 0.5 μm. **B)** Quantification of OPG staining intensity (*ImageJ*). **C)** ELISA performed on plasma samples from 80 MTC patients who were not treated (n=66) or treated with multi-tyrosine kinase inhibitors (vandetanib or cabozantanib) before blood collection (n=14). Circulating levels of OPG are shown according to *RET* M918T mutation status and bone metastases. **D)** Circulating levels of OPG are shown in patients with the M918T mutation who develop bone metastases and are either treated or untreated with tyrosine kinase inhibitors. **E)** Overall survival (OS) of MTC patients based on somatic *RET ^M918T^* mutation status and the presence of bone metastases. The median OS for no met/*RET*^WT^ and no met/*RET^M918T^* was not reached; and the median OS for met/*RET*^WT^ was 181 months; the median OS time for met/*RET^M918T^*was 179 months. (log-rank test, p=0.0089). **F)** Overall survival based on circulating OPG levels in plasma. OPG-low median OS was not reached; median OS for OPG-high was 181 months; **(log-rank test, *p*=0.0054)**.

In summary, these results indicate that circulating OPG in human plasma may provide a potential non-invasive approach to the diagnosis of MTC and determining the prognosis.

## DISCUSSION

This study shows that patient-derived MTC cells with somatic RET mutations (C634W or M918T) cause osteoblastic lesions in bone. Pharmacologic inhibition of RET suppresses tumor growth in bone and attenuates osteoblastic lesions. Mechanistically, activated RET mutation upregulates the secretion of OPG from MTC cells, leading to decreased osteoclastogenesis. Clinical studies of a large cohort of MTC patients showed that a high level of circulating OPG is associated with poor overall survival of MTC patients. Our work provides the first model of MTC tumor growth in bone, and out findings indicate that efficiently targeting RET activity and function attenuates tumor burden and skeletal lesions.

The non-hematopoietic cells of the bone marrow microenvironment include multipotent stromal cells, also called mesenchymal stem cells (MSCs), which have been defined in culture by their capacity to differentiate into OB, adipocytes, and chondrocytes [31–33]. Our *in vitro* studies showed that MTC cells induce the osteogenic differentiation of multipotent MSC by secreting growth factors and cytokines. However, osteoclastogenesis was strongly inhibited by MTC-CM cells. Among cytokines and growth factors secreted by MTC cells, OPG is the most prominent. OPG is a soluble RANK decoy receptor predominantly produced by OB that prevents OC formation. Overexpression of OPG leads to severe osteopetrosis in mice, and mice without OPG have bone loss [34–36]. Importantly, our data showed that RET knockdown or the pharmacologic inhibition of RET in MTC cells reduced OPG protein levels and secretion. OPG expression is reported to be regulated in OB by various cytokines (TNF-α, IL-1, TGFβ) and the Wnt/β-catenin signaling pathway [37]. RET is reported to interact with and activate β-catenin, which contributes to RET-mediated MTC tumor growth, invasion, and metastasis [38]. We speculate that RET activation of the Wnt/β−catenin pathway may lead to the upregulation of OPG in MTC cells.

MTC bone metastases have been seen in patients with mixed lytic and sclerotic bone metastases[1, 4]. Our data showed that MTC cells in the bone microenvironment produce a new type of woven bone, and such phenotypic changes in bone are influenced by the subtype of RET mutations in MTC cells. *RET^M918T^* mutations in MTC cells increased new bone formation in trabecular and cortical regions and caused porosity in the cortex.

Currently, bone metastases remain incurable, and therapies are mainly for the reduction of SRE’s as well as pain management. We found that selpercatinib, a RET-specific kinase inhibitor, is highly active against RET^M918T^ and RET^C634W^ mutant MTC tumors and inhibits tumor growth in bone. It is likely that selpercatinib inhibits MTC cell proliferation or survival and reduces MTC-induced abnormal bone formation. Additionally, ONC201, a small molecule, decreased MTC tumor growth in bone. We showed that ONC201 displayed no detectable side effects *in vivo* while reducing *in vivo* bone metastasis of RET-expressing MTC. Thus, inhibiting RET activity or protein levels can effectively target osteoblastic bone metastases due to MTC and reduce tumor growth in bone and morbidity and mortality in preclinical models.

Our results showed that patient-derived MTC cells cause distinct osteoblastic lesions in bone associated with RET mutations. Moreover, OPG is secreted by the MTC cells, and RET knockdown in these cells decreases OPG protein levels. Selpercatinib and pralsetinib inhibit OPG protein levels in MTC cells, and the OPG levels are higher in the primary tumor tissue of patients who develop bone metastases than in patients with no bone metastases or lymph node involvement. Thus, the presence of secreted OPG in human blood may provide a potential non-invasive approach for the diagnosis and prognosis of MTC patients in addition to the classic tumor biomarkers calcitonin and carcinoembryonic antigen. OPG may help detect MTC bone metastases early, even before symptoms arise or imaging shows clear evidence of metastases. This early detection allows for timely intervention and better management of the disease. OPG can track the progression of MTC bone metastases over time, as well as how well patients respond to treatment. OPG can be measured in the plasma, which is less invasive than traditional imaging methods, offering patients more convenient monitoring.

Studies have shown that RET mutations are amongst the most frequent cancer-inducing alterations in the germline of osteosarcoma patients, alongside mutations in tumor suppressor p53(TP53) and retinoblastoma protein 1 (RB1) and RET has been shown to be considered a driver of the disease. RET overexpression is associated with chemotherapeutic resistance to cisplatin and bortezomib and with increased stem-cell-like properties of osteosarcoma [39, 40]. Furthermore, elevated RET expression in breast tumors is associated with an increased risk of bone metastasis and poor overall survival [41–44]. Thus, our findings may have broader implications for osteoblastic lesions in other solid tumors.

### Limitations of the study

Although we show that MTC with activated RET mutations causes osteoblastic lesions by decreasing bone resorption via OPG, the mechanism by which RET regulates OPG remains to be determined. Furthermore, our understanding of how MTC alters the stromal composition and changes the intercellular communication networks in the bone microenvironment is incomplete. Addressing these questions will provide optimal therapy for treating MTC bone metastases.

## Resource availability

Any cell line generated by this project will be shared with the scientific community. The University of Texas MD Anderson Cancer Center will ensure that any legal restrictions on sharing these essential research tools are minimal.

## Lead contact

Further information and requests for resources and reagents should be directed to the lead contact, Dr. Theresa A. Guise.

## Material availability

Materials generated in this study will be made available upon request to the lead contact and with a materials transfer agreement with the University of Texas MD Anderson Cancer Center.

## Data and code availability

Raw data reported in this paper will be shared by the lead contact upon request. Any additional information required to reanalyze data reported in this paper is available from the lead contact upon request. No original code is reported in this paper.

## Acknowledgments

This work was supported by the Cancer Prevention Research Institute of Texas (Scholar of CPRIT Established Investigator Award RR190108: PI: TAG) and the Bone Disease Program of Texas (PI: R. B-Y). This work is also supported by the NIH/NCI under award number P30CA016672 (The University of Texas MD Anderson Cancer Center Support Grant) and the Small Animal Imaging Facility, the Bone Histomorphometry Core Laboratory, and the Research Histology Core Laboratory (UT MD Anderson Cancer Center). We thank Dr. Varun Prabhu for providing the ONC201 compound (*Chimerix Inc*, Durham, NC). The authors would like to thank Erica Goodoff of The University of Texas MD Anderson Cancer Center Medical Research Library for providing edits and proofing the manuscript.

## Author contributions

Conceptualization, R.B-Y. and T.A.G.; Methodology, R.B.Y.; Investigation, R.B-Y., G.M.P., J.L.K., T.T., D.A.R., L.M.C., J.M.M., XV, J.H.; Writing-Original Draft, R.B-Y.; Writing-Review & Editing, R.B-Y, G.M.P., J.L.K., T.T., M-C. H., S-H.L., M.I.H., S.I.S., T.A.G.; Funding Acquisition, R.B-Y. and T.A.G.; Resources, S.I.S; M.I.H.; Supervision, R.B-Y. and T.A.G.

## Declaration of interests

The authors declare no competing interests.

## MATERIALS AND METHODS

### Cell lines and primary OB

Human TT cells were purchased from ATCC (Manassas, VA). Human MZCRC1 cells were kindly provided by Dr. Alex Knuth (Switzerland) and were previously described [45–47]. RET mutations in MTC cell lines were verified by Sanger sequencing; TT cells harbor a codon 634 cysteine to tryptophan (C634W) exon 11 *RET* mutation, and MZCRC1 cells harbor a codon 918 methionine to threonine (M918T) exon 16 *RET* mutation. MTC cells were maintained in an F12/ Dulbecco’s modified Eagle medium with 10% fetal bovine serum (FBS), and HEK293T cells were cultured in Dulbecco’s modified Eagle medium with 10% FBS and supplemented with 10% serum. All cell lines tested negative for mycoplasma using the service provided by the Cell Cytology Core Facility at The University of Texas MD Anderson Cancer Center. The cells used for the experiments were between two and five passages from thawing. Primary OB were isolated from long mouse bones. After flushing bone marrow cells, trabecular bone pieces were dissected and placed into a phosphate-buffered saline (PBS) buffer. The bone pieces were washed three times in PBS and then further dissected into smaller bone pieces and incubated at 37°C for 20 minutes with shaking in 2 mg/ml collagenase type II (*Sigma Aldrich*, St. Louis, MO). The collagenase solution was then removed and discarded, and this procedure was repeated twice. The bone pieces were transferred into flasks containing Dulbecco’s modified Eagle medium. Culture medium was replaced 3 times per week. The OB growing from the bone fragments was passed 3 times before being used for the experiments. The same protocol was used for bones isolated from calvaria of newborn C57Bl/6 mice.

### Plasmids, lentiviral transduction, and RET knockdown

Lentiviral vectors (pLKO.1) containing RET-specific shRNAs were purchased from Sigma-Aldrich. Lentiviral RET-shRNA plasmids were co-transfected into HEK293T cells along with packaging (VPR8.9) and envelope (VSV-G) plasmids using X**-**tremeGENE (*Sigma Aldrich)* for two days. The virus particles containing RET-shRNA or control shRNA were used to infect TT and MZCRC1 cells. Transfected cells were selected in media containing 2 μg/mL puromycin (*Clontech,* San Jose, CA).

### Western blot analysis

Cells were lysed in NP40 lysis buffer (50 mM Tris-HCl [pH 7.4], 150 mM NaCl, 1% Np-40, 1 mM EDTA) containing phosphatase inhibitors (*Roche*) and protease inhibitors (*Pierce*, Waltham, MA). Proteins were transferred to PVDF membranes. After blocking in 5% bovine serum albumin in Tris-buffered saline with 0.05% Tween20, membranes were incubated with the primary antibody, followed by incubation with anti-rabbit (*Cell Signaling Technology*, Danvers, MA) or anti-mouse (*Cell Signaling Technology*) secondary antibodies conjugated with horseradish peroxidase. After washing with Tris buffer saline with 0.05% Tween-20, the bands were detected with a chemiluminescence reagent (*Thermo Fisher Scientific*, Waltham, MA). The primary antibodies used are as follows: antibodies against RET (*Cell Signaling Technology*), p-RET (*Santa Cruz Biotechnology, Inc,* Dallas, Texas), OPG (*Abcam,* Cambridge, UK), and vinculin as a housekeeping gene (*Sigma Aldrich*).

### Real-time qRT-PCR

Total RNA was isolated using the Direct-zol™ RNA Microprep kit (*Zymo Research™*, Tustin, CA). According to the manufacturer’s instructions, an aliquot of 200 ng of total RNA was subjected to reverse transcription with a SuperScript II RT-PCR kit (*Invitrogen*, Waltham, MA). Real-time qRT-PCR was performed using SYBR Green Master Mix (*Life Technologies*, Carlsbad, CA) or Taqman master mix with the QuantStudio 3 Real-Time PCR System (*Life Technologies*).

### *In vivo* mouse experiments, measurement of tumor burden, radiography, micro-CT, and bone histomorphometry

Animal experiments were approved by the University of Texas Institutional Animal Care and Use Committee (IACUC). Based on the known growth pattern of MTC cells in culture, 5 × 10^5^ tumor cells in 5-10 μL saline were administered directly through the femur while mice were maintained under anesthesia, and the contralateral femur was injected with PBS, which served as a control.

### Xray and DXA image analysis

Following tumor cell inoculation by weekly X-ray (*UltraFocus Faxitron*; Marlborough, MA), bone lesions were quantified in a blinded fashion using image analysis software (*OSTEO v20.1; Bioquant Imaging Analysis*, Nashville, TN). DXA scans were performed to quantify changes in whole body, tibial, femoral, and spinal bone mineral density, % body fat, and lean mass (*iNSiGHT, Osteosys*; London, ON) at study endpoint.

### Histology, TRAP staining, toluidine blue staining, and Von Kossa staining of the bone tissue

Mice were euthanized, and bones were collected, fixed in formalin for 2 d, washed, and transferred to 70% ethanol. Femurs were decalcified in 10% EDTA solution for 3 w, and embedded in paraffin, and the tissues were sectioned at 4 μm thickness. Slides were deparaffinized in xylene, rehydrated in gradients of ethanol, and incubated in PBS for 5 minutes. Sections were stained using hematoxylin, eosin, orange G, and phloxine. For TRAP staining, the slides were incubated with the staining buffer containing Fast Garn GBC solution, Naphthol AS-BI phosphoric acid solution, acetate solution, and tartrate solution in a 37C water bath protected from light for 1 hr. The slices were rinsed and counterstained with methyl green (*H-3402-500, Vector Laboratories*, Newark, CA), rinsed with water, and mounted with VectaMount Permanent Mounting Medium (*H-5000, Vector Laboratories*). The slides were stained with toluidine blue solution (*89640, Sigma*), dehydrated and cleared with xylene, and coverslipped with DMX hydrophobic adhesive. Undecalcified femurs were stained with Von Kossa stain (1% AgNO3, 5% Sodaformal, 5% NaThiosulfate). Bone histomorphometry was analyzed (*OSTEO v20.1, Bioquant Imaging Analysis*) on the region 150 μm from the distal growth plate and extending 1.3 mm into the bone compartment, keeping 150 μm distance from cortical walls. Within the measured bone compartment, the “Tissue Volume” measurement, the trabecular bone surface was traced, and static bone parameters were measured. OB were counted if they had the classic cuboidal shape with the characteristic dark nucleus and were in contact with the bone surface.

Lining cells were excluded because they were inactive. OC were multinucleated and were best visualized with TRAP enzymatic staining; they were measured only when in contact with the trabecular bone surface. The trabecular bone surface was also traced to provide structural data. OCy were quantified across the cortical surface of the bone due to the heightened OB differentiation and embedment within the bone. To assess tumor progression and bone quality, we determined the bone area, tumor area, OC #/mm at the tumor-bone interface, OC #/mm^2^, and OB #/mm^2^, OCy #/mm (*OSTEO v20.1, Bioquant Imaging Analysis*). Tumor progression and bone quality, bone area, tumor area, OC #/mm at the tumor bone interface, and OB #/mm were assessed by histomorphometry (*OSTEO v20.1, Bioquant Imaging Analysis*).

### Dynamic histomorphometry of bone

Dynamic histomorphometry was performed to determine the bone formation rate and phenotype. Calcein (*C0875, Sigma*,) was intraperitoneally injected twice (0.02 mg/gram of body weight) into mice (at 10 and 3 days prior to euthanasia) to obtain double-labeling of newly formed bones. The non-decalcified femurs were embedded in methyl methacrylate and sectioned at 5 μm thickness. Images were obtained using an inverted microscope.

### Immunohistochemical staining

Sections of mouse femora were deparaffinized in xylene and incubated in a gradient of ethanol. The slides were incubated in citrate buffer at 60 C overnight for antigen retrieval. After incubation in blocking buffer, slides were incubated with a primary rabbit antibody against OPG (1:200, *Abcam*) for 1 hr at room temperature and rinsed with PBS, followed by an incubation of 1 hr with the secondary biotinylated goat anti-rabbit antibody, and then the slides were treated with streptavidin-HRP for 30 minutes (*ABC Kit, Vector Laboratories*). For human tissues, after deparaffinization, slides were treated with boiling citrate buffer solution for 10 minutes using pressure cooker, and endogenous peroxidases were inactivated in a solution of 3% hydrogen peroxidase (*Sigma-Aldrich*) for 15 minutes and incubated for 1 hr with blocking buffer (*Avidin/Biotin Blocking Kit, Vector Laboratories*) followed by incubation with OPG antibody as described above. After three washes with PBS, slides were developed with 3,3’-diaminobenzidine (DAB) solution (*DAB Staining Kit, Vector Laboratories*). Counterstaining was performed with hematoxylin. Staining intensity was measured using Fiji imaging software (*ImageJ,* Bethesda, MD).

### *In vivo* micro-CT assessment of bone

For the selpercatinib studies, mouse femurs at week-3 of treatment were assessed using the VivaCT80 micro-CT scanner (*Scanco Medical*; Bruttisellen, Switzerland, voxel size=10 μm; integration time=300 ms). For the ONC201 *in vivo* studies, live mice were scanned using the SkyScan 1276 micro-CT scanner (*Bruker*, Billerica, MA, voxel size = 13 μm). Trabecular and cortical bone of distal femoral metaphysis were segmented and evaluated for 1 mm proximal to the growth plate. Cortical thickness (Ct.Th) and trabecular bone volume fraction (BV/TV), bone mineral density (BMD); average trabeculae number (Tb.N); average trabeculae thickness (Tb.Th) were two primary bone endpoints reported as well as 3D reconstructions for each sample.

### OC differentiation

Femurs and tibiae of C57BL/6 mice (male, 7 weeks old) were removed from mice after euthanasia. Bone marrow was extracted from the distal and proximal ends of the bone and filtered through a 70 µm cell strainer. Bone marrow cells were cultured in minimum essential medium (MEM) containing 10% serum for 72 hours and treated with 20 ng/mL of M-CSF. Adherent and non-adherent cells were collected and seeded in 96-well plates (20,000 cells/well) or 24-well plates (500,000 cells/well) and treated with 50 ng/ml of M-CSF (*315-02, Peprotech*, Rocky Hill, NJ) and 100 ng/mL of mouse soluble RANKL (*315-11, Peprotech*) for 5-6 days. The cells were cultured in the presence or absence of MTC-CM concentrated 100-fold using a 3K filter (*Millipore Sigma*, Burlington, MA), by centrifugation. According to the manufacturer instructions, TRAP staining identified the OC as multinucleated cells (*387A-1KT, Sigma-Aldrich*). Murine RAW264.7 cells were seeded in 24-well plates at the density of 5×10^5^ cells per well and cultured with 50 ng/ml of mouse soluble RANKL to induce OC differentiation in the presence or absence of MTC-CM concentrated 10-fold. TRAP staining for detecting mature OC was performed using a leukocyte acid phosphatase kit (*Sigma-Aldrich*) according to the manufacturer’s instructions.

### OB differentiation

Calvarial cells were isolated from newborn mice (C57BL6). Briefly, calvarial fragments without red hematopoietic regions were placed in a solution of collagenase type II and dispase (4 mg/ml each) in PBS. Calvaria pieces were incubated for 5 × 15-minute changes in total collagenase type II and dispase solution on a shaker at 37 ° C. Cells obtained from the first 15-minute incubation were discarded to exclude hematopoietic cells. Cells from the remaining four changes were pooled in tubes of fresh α-MEM with 20% FBS, centrifuged, and plated in α-MEM with 20% FBS in a 25 cm^2^ tissue culture flask. Fresh supplemented media was replaced the next day, followed by supplemented media change twice weekly. For producing and differentiating OB from long bones (tibiae and femora), the bone marrow and the bones were cut into small pieces and incubated with collagenase type II (1 mg/mL) for 90 minutes at 37C. After aspirating the digestion media, the bone pieces were cultured and passed three times before being used *in vitro*. Once pre-OB cells reached confluence (1–2 weeks), they were incubated in a medium containing 5 mmol·L^−1^ β-glycerophosphate and 100 μg·mL^−1^ ascorbic acid (*Sigma-Aldrich*) and replaced twice weekly. When the cells reached 90% confluence, osteogenic differentiation media containing 10mM β-glycerophosphate (*S3620, Selleck Chemicals;* Houston, TX), and 50 μg/mL ascorbic acid (*S3114, Selleck Chemicals*) was added, and cells were cultured for another 10-21 days. As indicated, the differentiation media was Sted with of MTC-CM cells concentrated 10-fold. Then, alkaline phosphatase staining was performed using a staining kit (*86R-1KT, Sigma*) according to the manufacturer instructions. A fluorometric assay was performed to measure the alkaline phosphatase activity. After osteogenic differentiation, the cells were washed twice with PBS and fixed with 4% formalin for 15 minutes and stained with 1% alizarin red-S (*A5533, Sigma-Aldrich*).

### ELISA

Mouse blood was collected through cardiac puncture immediately after euthanasia. The blood was collected into tubes with the anticoagulant EDTA and centrifugated at 2700 rpm in a refrigerated centrifuge for 15 minutes. The resulting supernatant was aliquoted and stored in an 80° C freezer. To detect human OPG an ELISA kit (*DY805, Bio-Techne*, Minneapolis, MN) OPG was used according to the manufacturer’s instructions. Human plasma was prepared from EDTA-treated blood within 4 hours of blood sample collection and frozen at -80°C.

### MTC patients and clinical data

All patients were treated at The University of Texas MD Anderson Cancer Center for “Sporadic MTC”. Patients gave written informed consent before collecting blood and tumor samples were collected under institutional review board-approved protocols. Participants had evidence of persistent/recurrent biochemical and/or structural disease. Tumor, node, metastasis (TNM) staging was based on the 7^th^ Edition of American Joint Committee on Cancer 7^th^ Edition criteria [48]. The date of diagnosis was defined as the date of pathologically confirmed MTC, either biopsy or initial surgery when the biopsy was indeterminate. Blood collection was performed concurrently with routine standard-of-care measurements, including blood calcitonin and carcinoembryonic antigen concentrations. *RET* germline mutation (exon 16) status was determined by Sanger sequencing.

### Statistical analysis

Results are presented as medians with range for continuous variables, and frequency with related percentages for the categorical variables. Statistical comparisons were conducted using Wilcox 2-sample tests for continuous variables, particularly OPG plasma levels, and the Fisher’s exact tests was employed to assess distribution differences in categorical variables. Statistical significance was indicated as follows: *, *p* <0.05; **, *p* <0.01; ***, *p*<0.001; and ****, *p* <0.0001.

Overall survival was defined as the elapsed time from blood collection to death or last follow-up date. Patients were censored at the last follow-up if death had not occurred. Univariate survival analysis was conducted using Kaplan-Meier estimates, with the log-rank test to compare the differences in overall survival curves between groups. The association between the clinicopathologic variables and overall survival was evaluated using the Cox regression model. *p*<0.05 was considered significant.

### Data availability

The data supporting this study’s findings are available within the article, its supplementary information files, and from the corresponding author upon request.

## Notes

### Competing Interest Statement

The authors have declared no competing interest.

