## Supplementary figures and images for "Activating *RET* Mutations Promotes Osteoblastic Bone Metastases in Medullary Thyroid Cancer"

### Supplemental Figure S1

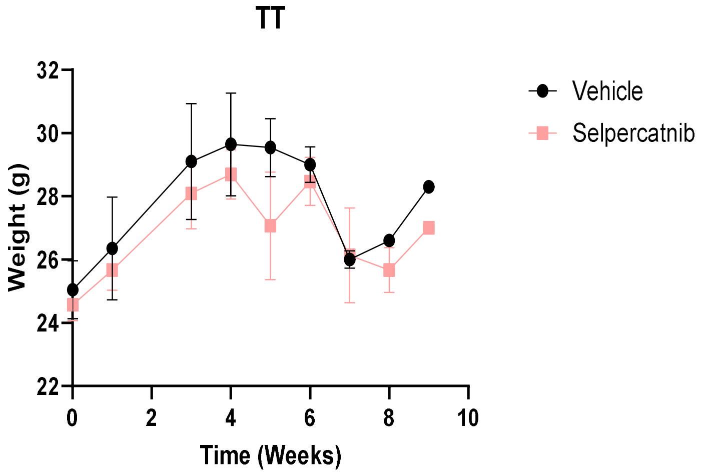


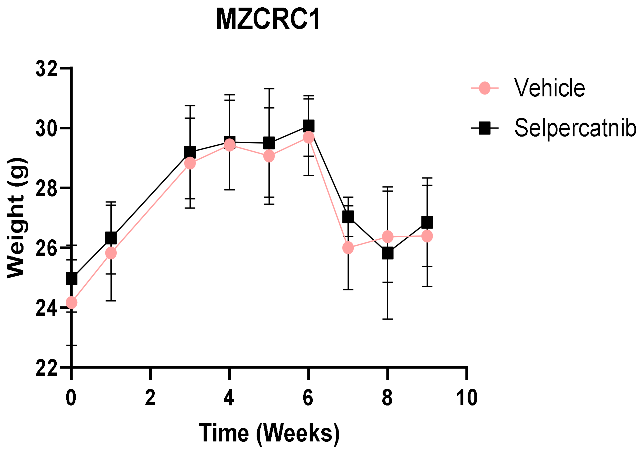


Supplemental Figure S1
