## Supplemental Table S1 for "Activating *RET* Mutations Promotes Osteoblastic Bone Metastases in Medullary Thyroid Cancer"

| **Table 1** Summary of Patients and Clinical Characteristics | | |
|---|---|---|
| Measure | Level | All (N=80) |
| RET mutations, n (%) | MTC-RET M918T | 52 (65) |
| | No RET-M918T | 27 (34) |
| | Not tested | 1 (1) |
| OPG | Median | 1256 |
| | Range | (650 – 3079) |
| OPG above 1000 pg/ml, n (%) | High OPG | 50 (62) |
| | Low OPG | 30 (38) |
| Stage, n (%) | Stage II | 4 (5) |
| | Stage III | 9 (11) |
| | Stage IV | 54 (68) |
| | Unstaged | 13 (16) |
| PT category, n (%) | T1 | 9 (11) |
| | T2 | 10 (12) |
| | T3 | 31 (39) |
| | T4 | 14 (18) |
| | TX | 16 (20) |
| PN Category, n (%) | N0 | 2 (3) |
| | N1/N1a | 4 (5) |
| | N1b | 60 (75) |
| | Nx | 2 (3) |
| | Missing | 12 (16) |
| M category, n (%) | M0 | 35 (44) |
| | M1 | 24 (30) |
| | MX | 21 (26) |
| Bone metastasis, n (%) | Bone Met | 43 (54) |
| | No Met | 37 (46) |
| Treatment, n (%) | Treated with drugs | 36 (45) |
| | No drugs | 44 (55) |
| Race, n (%) | White | 70 (88) |
| | Hispanic | 6 (8) |
| | Black | 3 (4) |
| | Asian | 1 (1) |
| Gender, n (%) | Female | 46 (58) |
| | Male | 34 (42) |
| Age | Number of subjects | 79 |
| | Median | 63 |
| | Range | (28 – 83) |
| Vital status, n (%) | Dead | 23 (29) |
| | Alive | 57 (71) |
| Follow-up months | Median | 124.4 |
| | Range | (37.8 – 501.1) |
