## Supplemental Table S2 for "Activating *RET* Mutations Promotes Osteoblastic Bone Metastases in Medullary Thyroid Cancer"

| **Table 2** Summary of Patients and Clinical Characteristics-by categorized OPG | | | | |
|---|---|---|---|---|
| Measure | Level | Low OPG (N=30) | High OPG (N=50) | P-value |
| RET mutations, n (%) | MTC-RET M918T | 19 (63) | 33 (66) | 0.58 |
| | No RET-M918T | 10 (33) | 17 (34) | |
| | Not tested | 1 (3) | 0 (0) | |
| Stage, n (%) | Stage II | 2 (7) | 2 (4) | 0.65 |
| | Stage III | 4 (13) | 5 (10) | |
| | Stage IV | 21 (70) | 33 (66) | |
| | Unstaged | 3 (10) | 10 (20) | |
| PT category, n (%) | T1 | 4 (13) | 5 (10) | 0.09 |
| | T2 | 2 (7) | 8 (16) | |
| | T3 | 12 (40) | 19 (38) | |
| | T4 | 9 (30) | 5 (10) | |
| | TX | 3 (10) | 13 (26) | |
| PN Category, n (%) | N0 | 2 (7) | 0 (0) | 0.28 |
| | N1/N1a | 2 (7) | 2 (5) | |
| | N1b | 24 (86) | 36 (90) | |
| | Nx | 0 (0) | 2 (5) | |
| | Missing | 2 | 10 | |
| M category, n (%) | M0 | 15 (50) | 20 (40) | 0.36 |
| | M1 | 10 (33) | 14 (28) | |
| | MX | 5 (17) | 16 (32) | |
| Bone metastasis, n (%) | Bone Met | 11 (37) | 32 (64) | 0.022 |
| | No Met | 19 (63) | 18 (36) | |
| Treatment, n (%) | Treated with drugs | 13 (43) | 23 (46) | 1.00 |
| | No drugs | 17 (57) | 27 (54) | |
| Race, n (%) | White | 27 (90) | 43 (86) | 0.35 |
| | Hispanic | 2 (7) | 4 (8) | |
| | Black | 0 (0) | 3 (6) | |
| | Asian | 1 (3) | 0 (0) | |
| Gender, n (%) | Female | 15 (50) | 31 (62) | 0.35 |
| | Male | 15 (50) | 19 (38) | |
| Age | Number of subjects | 30 | 49 | |
| | Median | 60 | 63 | 0.40 |
| | Range | (36 – 83) | (28 – 81) | |
| Vital status, n (%) | Dead | 5 (17) | 18 (36) | 0.08 |
| | Alive | 25 (83) | 32 (64) | |
| * P-value is from Wilcoxon two-sample test. | | | | |

| **Table 2 cont.** Summary of OPG plasma levels | | | | |
|---|---|---|---|---|
| Measure | Levels | Number of patients | Median (Range) | P-value* |
| Bone metastasis and RET mutations | No MET-WT | 13 | 1090 (670 – 1814) | 0.017 |
| | No MET-M918T | 24 | 1147.5 (689 – 2311) | |
| | MET-WT | 15 | 1425 (682 – 3079) | |
| | MET-M918T | 28 | 1653 (650 – 2887) | |
| Bone metastasis | Without | 37 | 1097 (670 – 2311) | 0.002 |
| | With | 43 | 1510 (650 – 3079) | |
| * P-value is from Wilcoxon two-sample test. | | | | |
