## Supplemental Table S3 for "Activating *RET* Mutations Promotes Osteoblastic Bone Metastases in Medullary Thyroid Cancer"

| **Table 3 Summary of Overall Survival Univariate Analysis** – months to death | | | | | | |
|---|---|---|---|---|---|---|
| | Overall Survival | | | | | |
| Measure | Level | Total#: #events | Median (95% CI) in months | p-value | Hazard Ratio (95% CI) | p-value |
| OS | | 80:23 | 417.6 (331.6 – NE) | | | |
| OPG status | Low | 30:5 | NE (331.63 – NE) | 0.040 | Ref (Low) | |
| | High | 50:18 | 417.6 (161.5 – 417.6) | | 2.73 (1.01 – 7.44) | 0.040 |
| | | | | | Ref (High) | |
| | | | | | 0.37 (0.13 – 0.99) | 0.040 |
| Bone metastasis | Without | 37:4 | NE (NE-NE) | 0.003 | Ref | |
| | With | 43:19 | 181.4 (155.7 – 417.6) | | 4.52 (1.53 – 13.34) | 0.003 |
| Bone metastasis and RET mutations | No Bone metastasis -WT | 12:0 | NE (NE-NE) | 0.017 | | |
| | No Bone metastasis -M918T | 24:4 | NE (NE-NE) | | | |
| | Bone metastasis - WT | 15:6 | 181.4 (90.6 – 417.6) | | | |
| | Bone metastasis - M918T | 28:13 | 331.6 (93.8 – NE) | | | |
| Bone metastasis (only among the RET M918T patients) | Without | 24:4 | NE (NE-NE) | 0.07 | Ref | |
| | With | 28:13 | 331.6 (93.8 – NE) | | 2.74 (0.89 – 8.42) | 0.07 |

| Table 3 cont. Summary of Overall Survival Univariate Analysis – Time to death or follow-up | | | | | | |
|---|---|---|---|---|---|---|
| | | Overall Survival | | | | |
| Measure | Level | Total#: #event | Median (95% CI) in months | p-value | Hazard Ratio (95% CI) | p-value |
| OS | | 80:23 | 417 (331–NE) | | | |
| OPG status | Low | 30:5 | NE (331 – NE) | 0.041 | Ref (Low) | |
| | High | 50:18 | 417 (161 – 417) | | 2.73 (1.00 – 7.43) | 0.041 |
| | | | | | Ref (High) | |
| | | | | | 0.37 (0.13 – 1.00) | 0.041 |
| Bone metastasis and RET mutations | No Bone metastasis -WT | 12:0 | NE (NE-NE) | 0.017 | Ref | |
| | No Bone metastasis -M918T | 24:4 | NE (NE-NE) | | | |
| | Bone metastasis - WT | 15:6 | 181 (90 – 417) | | | |
| | Bone metastasis - M918T | 28:13 | 331 (93 – NE) | | | |
